# Genetically encoded multimode reporter of adaptor protein 3 (AP-3) traffic in budding yeast

**DOI:** 10.1101/314286

**Authors:** Rachael L. Plemel, Greg Odorizzi, Alexey J. Merz

## Abstract

**SYNOPSIS:** The AP-3 (adaptor complex 3) mediates traffic from the late Golgi or early endosomes to late endosomal compartments. Here, a synthetic reporter is presented that allows convenient monitoring of AP-3 traffic, and direct screening or selection for mutants with defects in the pathway. The reporter can be assayed by fluorescence microscopy or in liquid or agar plate formats and is adaptable to high-throughput screening.

**SUMMARY:** AP-3 (adaptor complex 3) mediates traffic from the late Golgi or early endosomes to late endosomal compartments. In mammals, mutations in AP-3 cause Hermansky-Pudlak Syndrome type 2, cyclic neutropenias, and a form of epileptic encephalopathy. In budding yeast, AP-3 carries cargo directly from the *trans*-Golgi to the lysosomal vacuole. Despite the pathway’s importance and its discovery two decades ago, rapid screens and selections for AP-3 mutants have not been available. We now report GNSI, a synthetic, genetically encoded reporter that allows rapid plate-based assessment of AP-3 functional deficiency, using either chromogenic or growth phenotype readouts. This system identifies defects in both the formation and consumption of AP-3 carrier vesicles and is adaptable to high-throughput screening or selection in both plate array and liquid batch culture formats. Episomal and integrating plasmids encoding GNSI have been submitted to the Addgene repository.

## INTRODUCTION

Transport vesicle budding is mediated by coat protein complexes that shape the nascent carrier. Generally, coat complexes associate with adaptors that populate the transport vesicles with specific integral membrane cargo molecules. The heterotetrameric adaptor protein (AP) complexes 1 and 2 belong to a family of coat protein adaptors originally identified by means of their association with clathrin. AP-1 and AP-2 function with clathrin coat proteins to mediate vesicular transport from the trans-Golgi network and the plasma membrane, respectively. Several additional AP complexes have subsequently been identified.^1^ AP-3 functions at *trans*-Golgi and/or early endosomal compartments, sorting its cargo molecules into transport vesicles that fuse with lysosomes and lysosome-related organelles, a pathway parallel to but distinct from the classical vacuolar protein sorting (*VPS*) route. This functional assignment emerged from the discovery that mutations in AP-3 subunits correspond to many of the earliest known genetic mutations that affect pigmentation in laboratory strains of fruit flies and mice.^2–5^ In humans, mutations that disrupt AP-3 function cause an autosomal recessive disorder known as Hermansky-Pudlak Syndrome type 2, characterized by albinism, clotting deficiency, and a host of other abnormalities due to the malfunction of cell types that rely on lysosome-related organelles.^6, 7^

The identification of an AP-3-dependent protein transport pathway from trans-Golgi compartments to the lysosome-like vacuole in *Saccharomyces cerevisiae* highlighted the early evolution of AP-3 in eukaryotes.^8, 9^ The discovery of the yeast AP-3 pathway also led to expectations that yeast genetic screens could be used to characterize this transport pathway. Meeting these expectations, however, has proven difficult, because yeast use partially redundant pathways to deliver cargoes to the vacuole. Consequently, screens for genes that influence AP-3 function have relied on laborious assays of vacuolar ALP processing: either preparation of cell extracts followed by SDS-PAGE and immunoblotting for ALP, or ^[35]^S pulse-chase, followed by cell extract preparation, anti-ALP immunoprecipitation, SDS-PAGE, and autoradiography.^10, 11^ To date, only the immunoblot approach has been applied at near genomic scale. This required a multiyear effort, and essential genes were not surveyed.^10^ We now report a rapid and scalable phenotypic assay of AP-3 deficiency in yeast, based on a synthetic, genetically encoded reporter, GNSI. GNSI assesses AP-3 function using a colorimetric overlay, a positive selection for growth, or fluorescence microscopy, enabling convenient and scalable analyses of the AP-3 complex and its cofactors.

## RESULTS & DISCUSSION

Translational fusions to Suc2, the major invertase (INV) enzyme of budding yeast, have been a mainstay in studies of yeast endocytic trafficking. For example, carboxypeptidase Y (CPY) traffics from the late Golgi to the endosome, then to the vacuole, through a mechanism that requires endosome-Golgi recycling of Vps10, a transport receptor for CPY. The CPY itinerary defines the *VPS* pathway.^12^ Loss of Vps10, or *trans*-acting *vps* mutations that disrupt Vps10 cycling, result in exocytosis of CPY into the periplasmic space between the plasma membrane and cell wall. CPY-INV fusions, as well as fusions of vacuolar proteases A and C to INV, were used in pioneering screens that led to the identification of dozens of *VPS* genes.^13–15^ More recently, the v/R-SNARE Snc2 was fused to both green fluorescent protein (GFP) and invertase. The resulting reporter, GSS (here called GSI; Figure 1A) was used to assay the integrated levels of endocytosis and exocytosis in high-throughput screens of yeast gene deletion mutants.^16–18^

**Figure 1.**
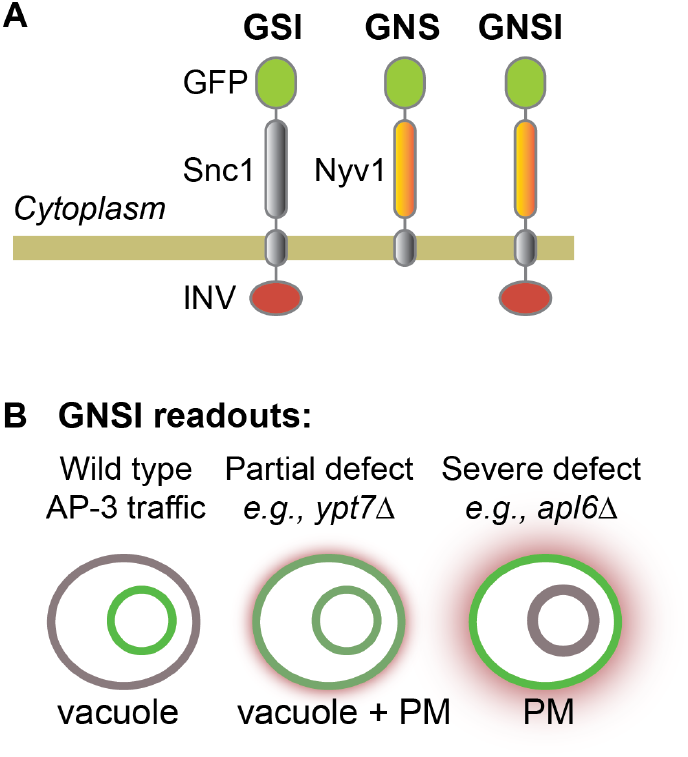
Optical and enzymatic reporters of endocytic recycling (GSI) and AP-3 (GNS, GNSI) traffic. **A.** Schematic diagram of the reporters. **B.** Known behavior of GNS, and predicted behavior of GNSI. In wild type cells, the GFP signal of GNS is localized almost exclusively to the vacuole, and the invertase (INV) reporter enzyme is degraded in the vacuole lumen. In AP-3 deficient cells, GNS is diverted to the plasma membrane (PM) and the invertase (INV) reporter enzyme is exposed to the extracellular space, where it can catalyze the conversion of a colorless, cell-impermeant substrate into a quantifiable colored product, depicted here as a halo.

The v/R-SNARE protein Nyv1 is normally targeted to the limiting membrane of the yeast vacuole through the AP-3 pathway.^19, 20^ Nyv1 sorting requires interaction between a YXXΦ-like motif (YGTI) in Nyv1’s N-terminal longin domain and Apm3, the AP-3 μ-chain.^20^ If AP-3 function is deficient, Nyv1 is mis-routed into the endosomal system. Nyv1 then enters the multivesicular body (MVB), and is destroyed in the lumen of the lysosomal vacuole.^20^

Previously, Reggiori *et al.*^19^ studied a chimera called GNS (Figure 1A). In GNS, the GFP-marked Nyv1 cytoplasmic domain is fused to the transmembrane anchor of the v/R-SNARE, Snc1. In wild type cells GNS, like Nyv1, is targeted to the vacuole limiting membrane. In cells defective for AP-3 function, however, GNS is diverted not to the MVB but instead to the plasma membrane, where it is trapped. GNS has since been used as an optical reporter of AP-3– mediated traffic.^20–22^ Based on these results, we reasoned that fusion of invertase to the exoplasmic C-terminus of GNS would yield a reporter (GNSI; Figure 1A) that could be used to monitor AP-3 traffic, using plate-based or liquid culture systems (Figure 1B). We constructed the GNSI chimera and expressed it from a single-copy plasmid under the constitutive *CYC1* promoter. We then evaluated GNSI as an AP-3 reporter using three readouts.

First, we asked whether GNSI faithfully recapitulates the subcellular localization of the original GNS reporter.^19^ As shown in Figure 2, it does so. In wild type cells, GNSI is targeted efficiently to the vacuole limiting membrane. In cells lacking AP-3 (*apl6*Δ) or in cells defective for terminal docking and fusion of AP-3 vesicles at the vacuolar target membrane (*vam3^tsf^*) GNSI accumulated largely on the plasma membrane. In cells with a less penetrant AP-3 deficiency (*ypt7*Δ), plasma membrane localization was weaker and observed only in a subset of the cells. Because AP-3 mutant cells exhibited some GNSI staining on the vacuole as well as the plasma membrane, we asked whether this localization depends on endosomal trafficking (Supplementary Figure S1). Single and double mutants were constructed in *apm3* (AP-3 μ subunit), *pep12* (endosomal Qa-SNARE), and *vps4* (ATPase required for ESCRT-mediated cargo sequestration in multivesicular bodies). As expected, the *pep12*Δ and *vps4*Δ mutants exhibited VPS class D and E vacuole morphology, respectively, but efficiently sorted GNSI to the vacuole (Figure S1), with little or no signal detected on the plasma membrane. The double *apm3*Δ *pep12*Δ mutant had a “shattered” VPS class C vacuolar morphology, with FM4-64 staining visible in the cytoplasm but no discernable vacuolar structures, while the *apm3*Δ *vps4*Δ mutant displayed a strong class B morphology, with fragmented vacuoles. In both double mutants GNSI accumulated on the plasma membranes of most cells, and was also visible on intracellular structures, consistent with leaky transport to the vacuole. In experiments using high-copy (rather than single-copy) vectors, or promoters stronger than *CYC1pr*, we noted that GNSI tended to accumulate in the endoplasmic reticulum (unpublished data). Thus, overexpression is best avoided with GNSI. With this caveat, the subcellular localization of GNSI closely mirrors the localization of the original GNS reporter.

**Figure 2.**
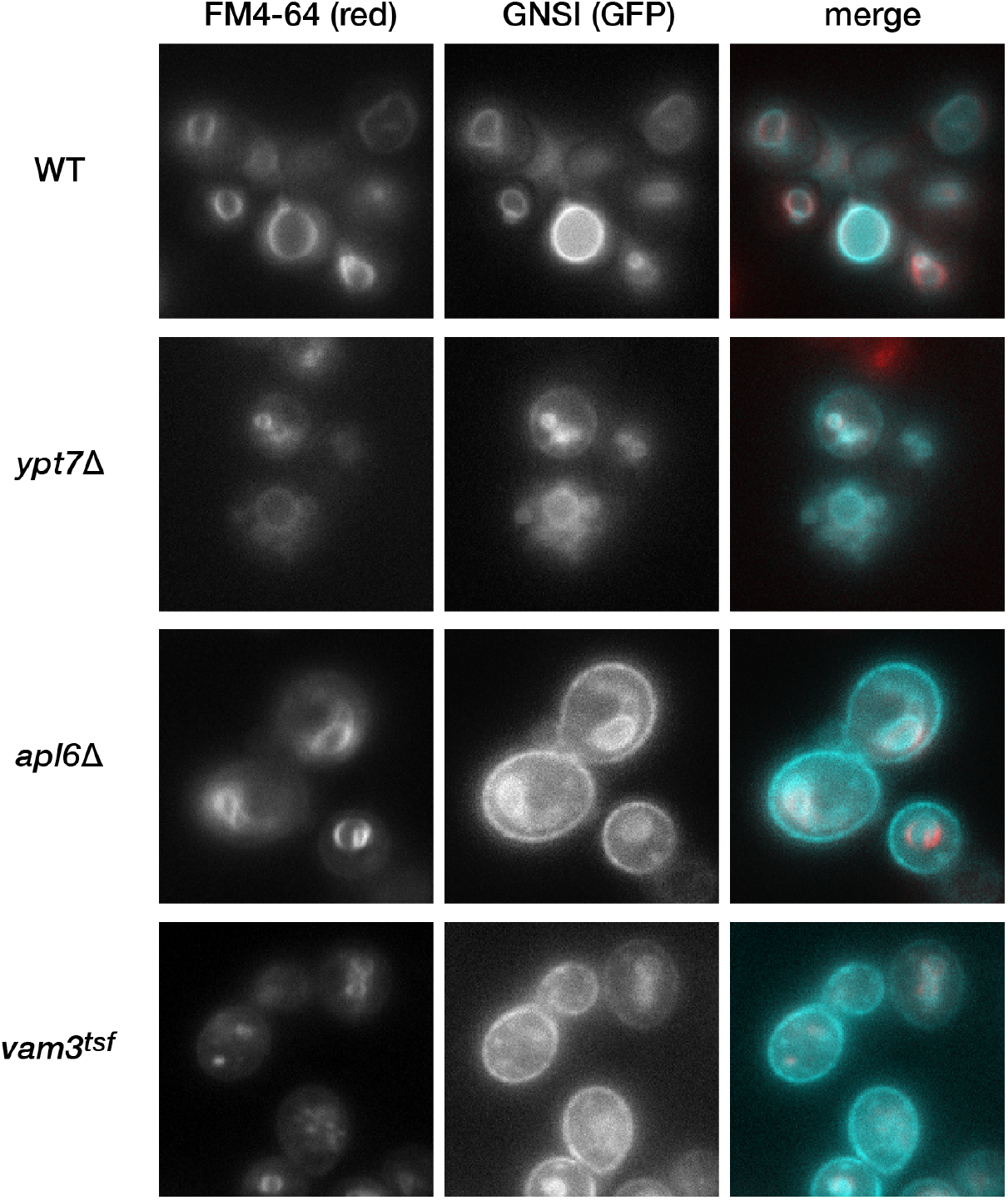
Subcellular localization of the GNSI probe. Cells were grown to mid-log phase at 30° C, and vacuoles were pulse-chase labeled with the red styryl dye FM4-64. Wide-field fluorescence micrographs are shown. In the color merge panels, the GFP signal is false-colored cyan. All strains here are in the SEY6210 genetic background.

Next, we tested whether a chromogenic assay that measures glucose liberation by invertase^23^ could be used as a plate-based assay of AP-3 activity. As shown in Figure 3, colored invertase reaction product strongly accumulated in strains defective for AP-3 traffic. As with the microscopy readout, strains previously reported to exhibit more severe trafficking defects using proteolytic processing of ALP as a readout also yielded stronger signals with the GNSI reporter.

**Figure 3.**
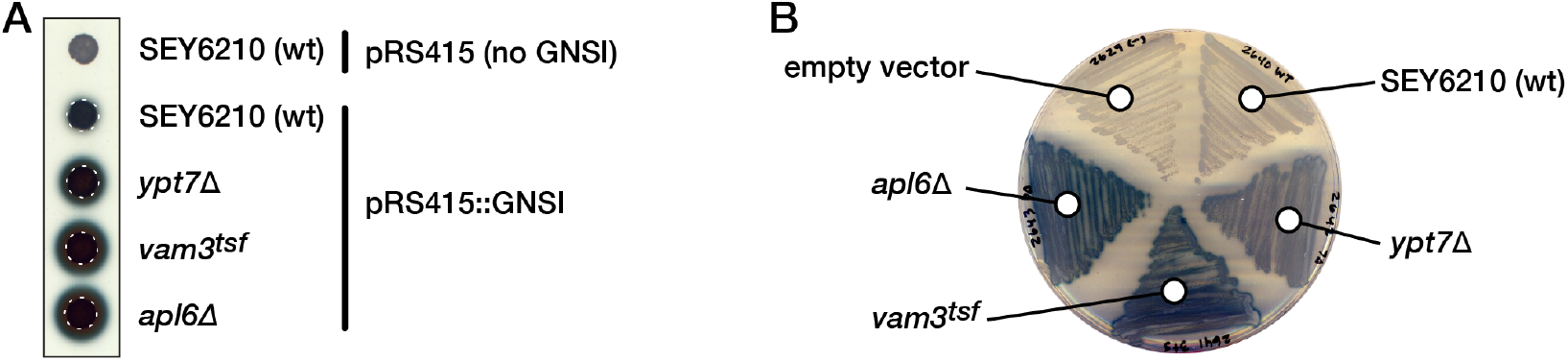
Agar overlay assay of defects in AP-3 trafficking. Liberation of glucose by invertase exposed to the extracellular (periplasmic) space was detected as described in Methods. Dark precipitate indicates the presence of glucose. **A.** Colonies grown on 2% fructose solid media following inoculation with a pin manifold (48-well spacing) are assayed using the invertase overlay technique. Dashed circles denote the colony diameters. **B.** Streaked sectors of a 2% fructose agar plate are assayed. All strains are in the SEY6210 genetic background.

Proteolytic processing of ALP requires not only that ALP be targeted to the vacuole, but that vacuolar proteases be both correctly localized and active in the vacuole lumen. Thus, accumulation of unprocessed ALP precursor (proALP) does not necessarily indicate a defect in the sorting or vacuolar delivery of AP-3 cargoes. To assess whether vacuolar hydrolase activity is needed for GNSI to report wild type AP-3 function, we assayed a *pep4*Δ null mutant. *PEP4* encodes vacuolar protease A, a master protease required for the activation of several other vacuolar hydrolases ^24^, including protease B (Prb1). Together, proteases A and B are responsible for maturation of proALP.^25^ In a *pep4*Δ mutant, ALP maturation is severely impaired (Figure 4). This result does not reflect an error in AP-3, sorting, but rather a defect in proteolysis at the vacuolar destination compartment. In contrast, the GNSI overlay assay reports wild type AP-3 sorting in a *pep4*Δ mutant (Figure 4B). Thus, GNSI not only reports bona fide defects in AP-3 traffic, but also evades a major class of false positive AP-3 sorting defects reported by assays of ALP maturation.

**Figure 4.**
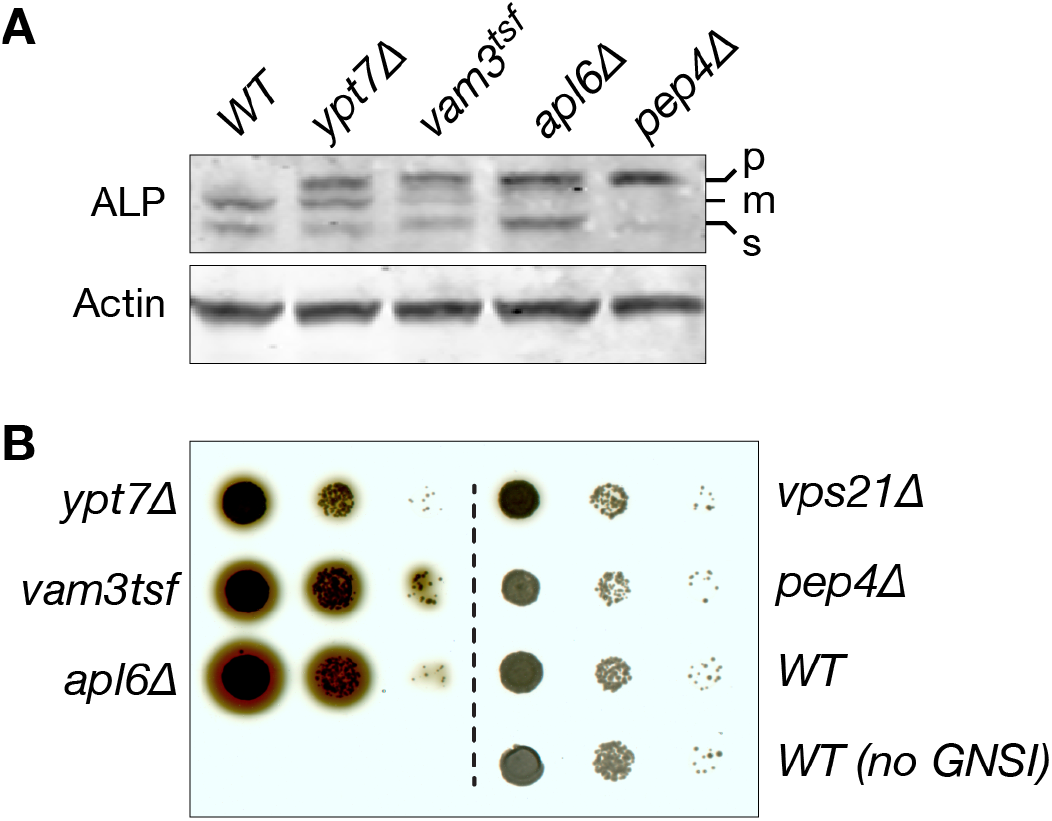
GNSI reports AP-3 trafficking defects but does not report vacuolar protease defects. **A.** Whole cell lysates were prepared from the indicated mutant strains, separated using SDS-PAGE, and immunoblotted with monoclonal anti-ALP (top) or anti-Actin (bottom). **B.** GNSI invertase overlay assays were performed on the indicated strains. Each strain was grown in liquid media to mid-log phase, then 1x, 1/10x, and 1/100x dilutions were inoculated onto 2% fructose solid media with a pin manifold, and the resulting colonies assayed by invertase overlay.

A limitation of the plasmid-borne GNSI reporter demonstrated above is that it can be used only in strains with the *suc2* genotype (e.g., SEY6210).^15^ However, by replacing the chromosomal *SUC2* locus with GNSI in cells also bearing the “magic marker” haploid selection locus^26^, it is possible to efficiently introduce *suc2Δ*::*GNSI* into any compatible mutant background through scalable mating and sporulation procedures.^27^ We therefore constructed the integrating vector p*SUC2*::*GNSI(NrsR)* to facilitate chromosomal replacement of *SUC2* with *GNSI*. As a proof of principle, we used this system to test mini-arrays of mutants assessed by ALP processing in two previous screens. First, we assayed a panel of missense alleles of *VPS33* (Figure 5). Vps33 is the SM (Sec1/mammalian UNC18) subunit of the CORVET and HOPS multi-subunit tethering complexes. In our previous studies these mutants exhibited heterogenous ALP processing, CPY processing, Zn^2+^ sensitivity, and vacuolar morphology phenotypes.^28^ Results using GNSI expressed from integrated suc2::*GNSI* tracked well with our previous results with ALP, indicating that the ALP maturation defects we had previously characterized do indeed reflect failures in AP-3 traffic, and not merely impaired vacuolar protease activity or localization.

**Figure 5.**
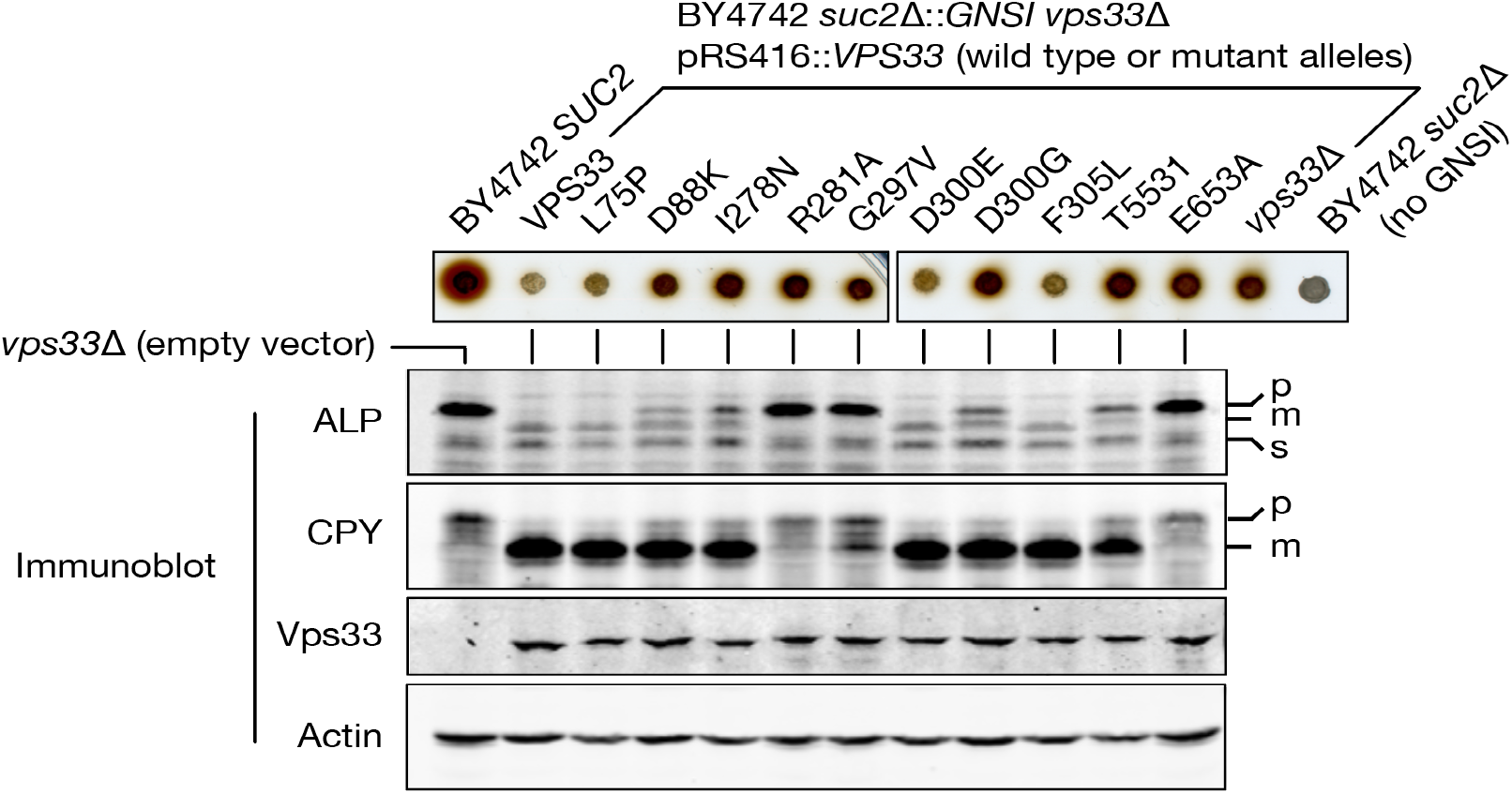
GNSI assay of AP-3 sorting defects conferred by *vps33* missense mutants. BY4742 *vps33Δ GNSI*::*suc2Δ* strains were transformed with the indicated pRS416::X plasmids and grown in liquid media. 1x, 1/10x, and 1/100x dilutions were inoculated onto 2% fructose agar media with a pin manifold, and the resulting colonies were assayed by invertase overlay. Isogenic strains lacking GNSI were previously characterized by immunoblot.^28^ The blots from that study are reproduced in this figure to allow direct comparison with the GNSI overlay results.

Next, we used the Magic Marker method to assess a subset of mutations that had been identified as candidate AP-3 trafficking factors in a large-scale screen for AP-3 trafficking defects (Figure 6).^10^ Mutants exhibiting defects in retrograde traffic from the endosome to the Golgi (*vps5*Δ, *vps29*Δ) also had mild AP-3 trafficking defects as assayed both by ALP maturation and GNSI. Thus, there are *bona fide* AP-3 sorting defects in these mutants, independent of defects in vacuolar protease activity. In contrast, GNSI reported wild type AP-3 traffic in *vps1*Δ, *vps8*Δ, *vps17*Δ, or *vps35*Δ mutant cells. The previous ALP maturation-based identification of these genes may reflect defects in vacuolar protease localization or activity, rather than defects in AP-3 traffic. Alternatively, the *vps17*Δ, or *vps35*Δ mutant cells assayed here might carry extragenic mutations that suppress a defect in AP-3 traffic. Overall, these results indicate that GNSI is suitable for large-scale screening using automated mating and sporulation procedures.^17, 26^

**Figure 6.**
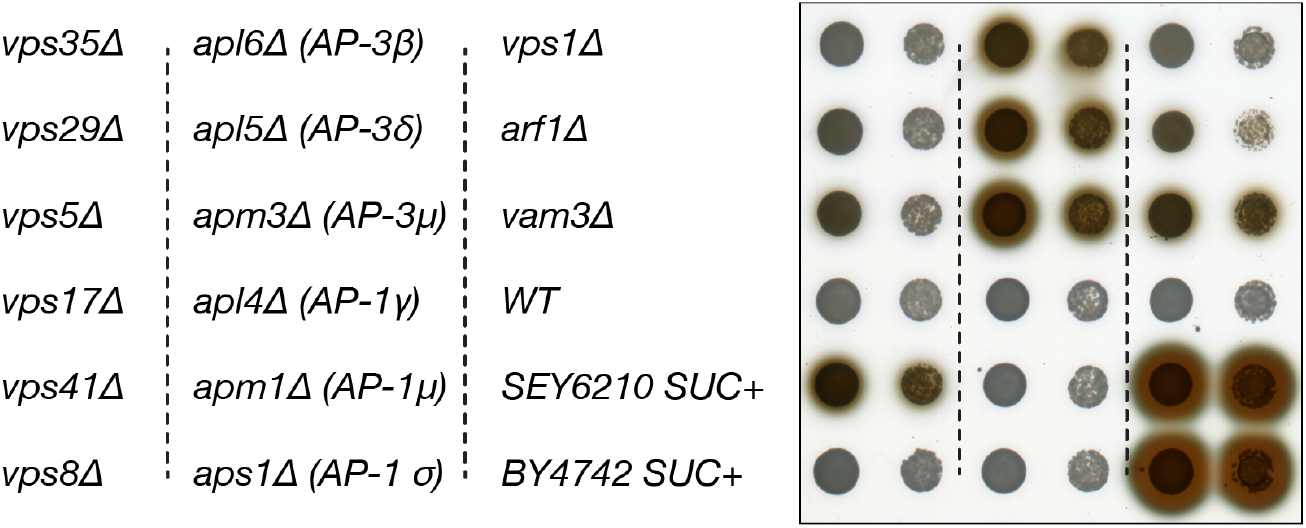
AP-3 traffic in mutant strains from the yeast gene deletion collection. Strains carrying the indicated null mutations were generated by mating to the Magic Marker *suc2Δ*::*GNSI* strain, followed by the selection of haploid spores carrying *GNSI* and the indicated null alleles. The resulting strains were grown in liquid media, then 1x and 1/10x dilutions were inoculated onto 2% fructose agar media with a pin manifold, and the plate was assayed by invertase overlay.

Finally, we asked if GNSI could be used in growth selections. Mutants lacking extracellular invertase exhibit growth defects when sucrose rather than dextrose is the primary carbon source. Thus, we tested whether *suc2*Δ cells with AP-3 deficiency would exhibit enhanced growth on sucrose plates containing the pH indicator dye bromocresol purple. As shown in Figure 7, cells expressing GNSI grew into larger colonies, and more strongly acidified the medium, when mutations compromising AP-3 traffic were present. As expected, the stringency of this growth selection was enhanced when oxidative respiration was blocked with the uncoupling toxin antimycin A (Figure 7). Thus, the GNSI reporter facilitates positive selection, with controllable stringency, of mutants defective in AP-3 traffic. Although we have not in this report quantified AP-3 sorting defects, approaches for quantifying surface-localized invertase reporters have been presented. These methods employ liquid culture or plate-based formats, with the latter being particularly amenable to high-throughput screening.^20, 23^

**Figure 7.**
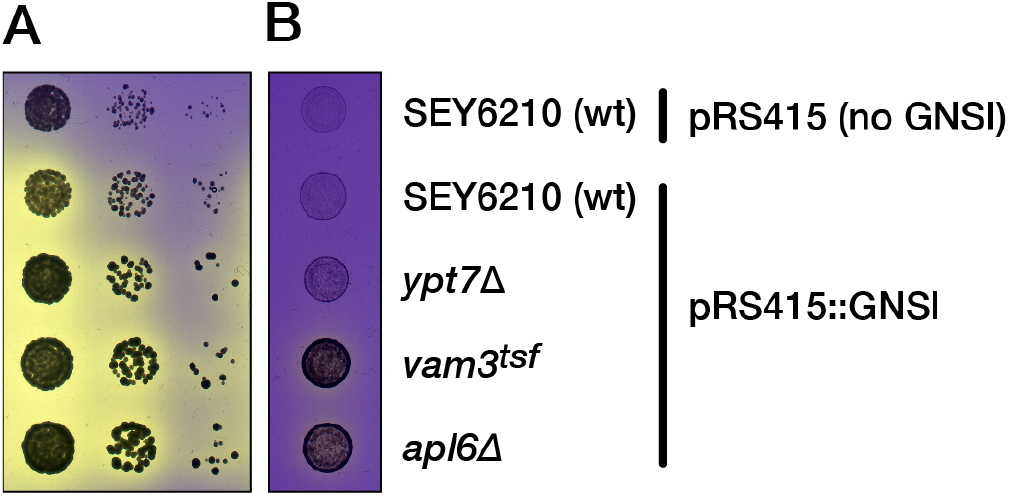
Growth phenotypes on sucrose indicator media. **A**. Ten-fold serial dilutions were inoculated with a pin manifold onto 2% sucrose agar media containing bromocresol purple. **B**. Cells were inoculated as in the first column of A, but onto media containing 10 μg/mL antimycin A. All strains are *suc2* mutants.

In summary, GNSI is a convenient and extremely versatile reporter of AP-3 traffic. Its use does not require laborious preparation and analysis of cell extracts or metabolic labeling. This system facilitates rapid and high-throughput analyses of AP-3 structure and function as well as genome-scale primary and secondary screening of potential AP-3 co-factors. Importantly, GNSI readily detects defects not only in AP-3 vesicle formation, but in the docking and fusion of AP-3 vesicles at the vacuolar target membrane. The AP-3 and VPS pathways, homotypic vacuole fusion, and autophagosome fusion at the vacuole all employ a common set of docking and fusion factors: Rab7/Ypt7, its GEF Ccz1/Mon1, its effector, the HOPS tethering complex, and a common set of SNAREs. To our knowledge GNSI is the first reporter that allows *direct positive selection* of mutants defective for docking and fusion in *any* of these pathways. This capability will in turn allow high-density deep-scan mutagenesis (DMS) structure-function studies^29^ of specific AP-3 transport factors, including Vps33 and the AP-3 complex itself. Similarly, we are using GNSI reporters to carry out genome-scale screens for new mutants defective in AP-3 transport.

## METHODS

### Cloning

pGNS416, encoding GFP-Nyv1-Snc1TMD^19^ was a gift from F. Reggiori. pHB4, encoding GSI (in earlier work called GSS)^18^ was a gift from E. Conibear. To construct p*CYC1pr-GNSI*(*LEU2*), the yeast vector pRS415 was linearized using XhoI and SacI. Using a 3-piece Gibson assembly procedure, three purified PCR products were cloned into the linearized vector: the *CYC1* promoter (290bp) was cloned from yeast genomic DNA, the *GNS* cassette was amplified from pGNS416, and the invertase cassette (including the *SNC1* TMD, *SUC2* ORF, and *SUC2* terminator) was amplified from pHB4. The resulting plasmid was amplified in *E. coli*, mapped using PCR, and fully sequenced using dideoxy chain termination chemistry. The GNSI integrating plasmid p*SUC2::GNSI(NrsR)* was constructed by digesting p*CYC1pr-GNSI*(*LEU2*) with ApaI (NEB), then using Gibson assembly to insert the *SUC2* promoter and *NrsR* cassette which were PCR amplified from pHB4. The resulting plasmid was amplified in *E. coli*, mapped using PCR, and fully sequenced using dideoxy chain termination chemistry. Plasmid DNA sequences and maps were designed using the SnapGene package (GSL Biotech). p*CYC1pr-GNSI*(*LEU2*) and p*SUC2::GNSI(NrsR)* plasmid DNA, maps, and full DNA sequence files, are available from the Addgene repository (plasmids #111450 and #121537, respectively).

### Yeast strains and media

#### Yeast strains

All yeast strains share the SEY6210 or S288C genetic background. The Magic Marker *GNSI* integrated strain was cloned by first digesting the p*SUC2::GNSI(NrsR*) plasmid with NotI, SacI, and EcoRV using NEB enzymes and CutSmart buffer. The EcoRV site is located within the *LEU2* coding sequence, and we found that digesting with this enzyme reduced the possibility of incorrect chromosomal integration. The digested plasmid DNA was then transformed into the yeast Magic Marker strain Y8205 (MAT**α**) from the Boone Lab and selected on YPD containing Nourseothricin. The *SUC2*::*GNSI* integration was mapped using PCR and verified by Sanger sequencing. The Magic Marker *GNSI* strain was then mated with a selection of BY4741 deletion collection strains (KanR) and the subsequent diploid cells were sporulated on ESM plates. Haploid spores containing the gene deletion and GNSI integrated cassette were selected using SD/MSG -His -Lys -Arg Canavanine + Thialysine + G418 + Nourseothricin plates.

#### Invertase agar overlay assay

The yeast strains were grown as liquid cultures in synthetic complete (SC) –Leu at 30° C to mid-log phase, then diluted to an OD^600^ of 0.5. Serial dilutions were pinned using a 48-pin manifold on (SC) –Leu, +2% fructose plates. The fructose plates were grown for one day at 30°, then the invertase agar overlay assay was done as described ^23^. Reactions were developed for 15 minutes before being photographed. The strains in Figure 5 were assayed under the same conditions, but were grown in (SC) –Ura and plated on (SC) –Ura, +2% fructose plates. The integrated GNSI strains in Figure 6 were grown in (SC) and plated on (SC) +2% fructose plates.

#### Sucrose indicator plate assay

The yeast strains were grown as liquid cultures in synthetic complete (SC)-Leu at 30° C to mid-log phase, then diluted to an OD^600^ of 0.5. Serial dilutions were pinned using a 48-pin manifold onto 2% sucrose indicator solid media ^23^ containing bromocresol purple (Thermo-Fisher Scientific), in the absence or presence of 10 μg/mL antimycin A (Fisher Scientific). The BP (bromocresol purple) indicator plates were grown at 30° C for two days before imaging.

### Microscopy

FM4-64 pulse chase was done as described ^30^ with a 20 minute exposure to the dye followed by a 30 minute chase in SC media without the dye. Epifluorescence micrographs were acquired using an Olympus IX71 microscope equipped with an Andor 885 camera, an intermediate 1.5X magnification lens, and a 60× 1.4NA objective.

## ACKNOWLEDGEMENTS

This work was supported by a seed grant from the University of Washington to AJM, and by grants to AJM and GO from NIH: GM077349, GM111335, and GM130644. We thank F. Reggiori and E. Conibear for sharing plasmids and D. Nickerson for illuminating discussions. The authors attest that they have no conflicts of interest.

---

**Supplementary Figure S1.**
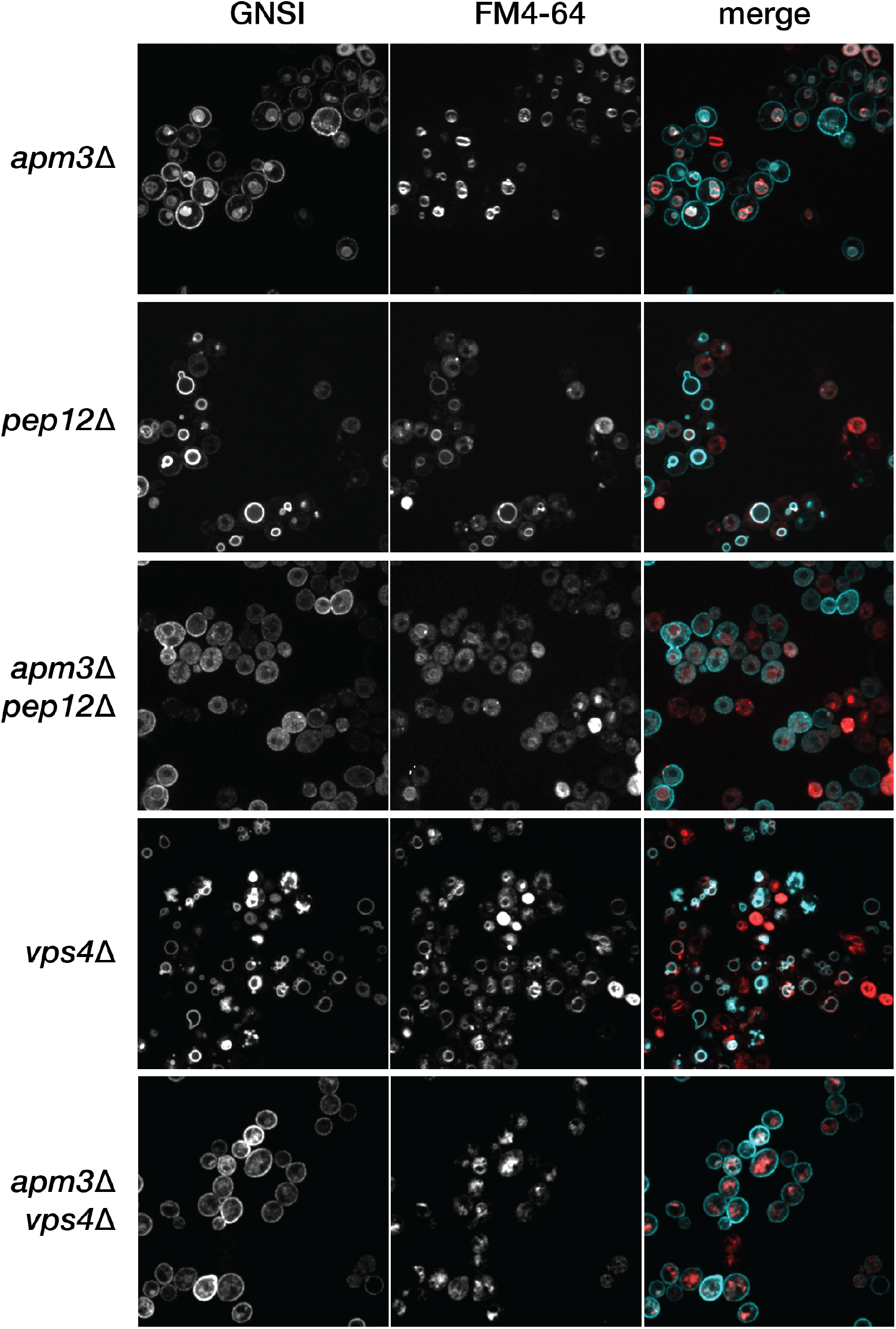
Subcellular localization of the GNSI probe in double mutants defective for both AP-3 and endosomal traffic. The indicated mutants were grown to mid-log phase at 30° C, and vacuoles were pulse-chase labeled with the red styryl dye FM4-64. Wide-field fluorescence micrographs are shown. In the color merge panels, the GFP signal is false-colored cyan. All strains here are in the SEY6210 genetic background.

